# Caprine humoral response to Burkholderia pseudomallei antigens during acute melioidosis from aerosol exposure

**DOI:** 10.1101/420075

**Authors:** Jinhee Yi, Mukoma F. Simpanya, Erik W. Settles, Austin B. Shannon, Karen Hernandez, Lauren Pristo, Mitchell E. Keener, Heidie Hornstra, Joseph D. Busch, Carl Soffler, Paul J. Brett, Bart J. Currie, Richard A. Bowen, Apichai Tuanyok, Paul Keim

## Abstract

*Burkholderia pseudomallei* causes melioidosis, a common source of pneumonia and sepsis in Southeast Asia and Northern Australia, that results in high mortality rates. A caprine melioidosis model of aerosol infection that leads to a systemic infection has the potential to characterize the humoral immune response. This could help identify immunogenic proteins for new diagnostics and vaccine candidates. Outbred goats may more accurately mimic human infection, in contrast to the inbred mouse models used to date. *B. pseudomallei* infection was delivered as an intratracheal aerosol. Antigenic protein profiling was generated from the infecting strain MSHR511. Humoral immune responses were analyzed by ELISA and western blot, and the antigenic proteins were identified by mass spectrometry. Throughout the course of the infection the assay results demonstrated a much greater humoral response with IgG antibodies, in both breadth and quantity, compared to IgM antibodies. Pre-infection sera showed multiple immunogenic proteins already reactive for IgG (7-20) and IgM (0-12) in most of the goats despite no previous exposure to *B. pseudomallei*. After infection, the number of IgG reactive proteins showed a marked increase as the disease progressed. Early stage infection (day 7) showed immune reaction to chaperone proteins (GroEL, EF-Tu, and DnaK). These three proteins were detected in all serum samples after infection, with GroEL immunogenically dominant. Seven common reactive antigens were selected for further analysis using ELISA. The heat shock protein GroEL1 elicited the strongest goat antibody immune response compared to the other six antigens. Most of the six antigens showed the peak IgM reactivity at day 14, whereas the IgG reactivity increased further as the disease progressed. An overall MSHR511 proteomic comparison between the goat model and human sera showed that many immune reactive proteins are common between humans and goats with melioidosis.

**Author Summary:** *B. pseudomallei* infection, the causative agent of melioidosis, results in severe disseminated or localized infections. A systemic study of the humoral immune response to *B. pseudomallei* infection using the *B. pseudomallei* aerosol caprine model would help understand the detectable antigenic proteins as the infection progresses. To study the immune response, IgG and IgM antibody responses to whole cell lysate proteins were identified and analyzed. Antigenic carbohydrates were also studied. From the results, this study suggests that the caprine humoral immune response to aerosolized *B. pseudomallei* has similarities to human melioidosis and may facilitate the analysis of the temporal antibody responses. In addition, commonly detected immunogenic proteins may be used as biomarkers for the future point of care (POC) diagnostics.

## Introduction

*Burkholderia pseudomallei* is a Gram-negative, non-spore forming, aerobic, and motile bacillus [1] and the etiological agent of melioidosis. This disease has emerged as a significant public health threat in Southeast Asia and Northern Australia [2]. Both *B. pseudomallei* and its close relative, *B. mallei*, the cause of glanders, are classified by the Centers for Disease Control and Prevention as category B bioterrorism agents [3]. In Thailand, *B. pseudomallei* is widely distributed in water and wet soils, such as rice paddies [4, 5]. In the two highly endemic regions of Northern Australia and Northeast Thailand, *B. pseudomallei* is responsible for a melioidosis fatality rate of around 10% and 40%, respectively [2, 5, 6]. There is also emerging evidence that melioidosis is endemic in Central and South America [7, 8], southern regions of China [9, 10] and India [11, 12]. The global distribution of *B. pseudomallei* endemicity has been linked to anthropogenic dispersal, both ancient and more recent [13]. Furthermore increased travel [14], and soldiers returning from endemic countries have led to many cases in non-endemic regions such as the USA and Europe [14, 15]. Limmathurotsalkul *et al* reported an evidence-based estimate of *B. pseudomallei* global distribution across the tropics; 46 countries were identified as suitable for melioidosis and with environmental suitability for the persistence [16]. Detection of cases outside endemic countries is also now helped by an increased awareness of melioidosis by clinicians worldwide [2]. The main routes of *B. pseudomallei* infection are dermal inoculation, inhalation and ingestion [17, 18]. Melioidosis clinical presentation spans from pneumonia (50% of all cases) [18, 19], and sepsis often leading to septic shock often with multiple abscesses in internal organs such as spleen, liver, kidney and prostate [20], to chronic abscesses in the skin without sepsis [18, 21].

Melioidosis is successfully treatable if diagnosed early and correctly. However, confirmed evidence of melioidosis infection in a patient currently relies on isolation and identification of *B. pseudomallei* from culture, often requiring use of selective medium [22, 23]. The bacteriological method takes a minimum of 24-48 hours, making it too slow to guide early treatment, which is particularly problematic for severe sepsis with its high mortality rate [2, 23-25]. To improve the diagnosis of melioidosis, a number of techniques have been attempted, such as antigen detection in specimens, antibody detection, molecular and rapid culture techniques [2]. A number of serological tests for antibody detection have also been developed for possible early diagnosis of melioidosis, viz. indirect hemagglutination (IHA), complement fixation (CF), immunofluorescence (IFA) and enzyme linked immunosorbent assay (ELISA) [2, 23, 26]. However, there are a number of problems associated with the interpretation of the results and sometimes the tests give false positive results [27]. Therefore, a sensitive, specific and rapid test is still needed for diagnosis of melioidosis, especially for those presenting with severe sepsis [6, 23]. Recombinant proteins as diagnostic antigens for serology may offer advantages over IHA, which is currently the only serological assay which used. The current IHA tests have low sensitivity early in infection plus poor specificity in endemic regions due to prior background exposure; i.e. false positives are not uncommon [2].

In the last 10 years, melioidosis and *B. pseudomallei* have attracted increased global attention and the related research has likewise increased considerably. However, there are large gaps in our knowledge of the pathogenesis of *B. pseudomallei* infection and the host immune response [6]. The comparative studies of an animal model and human infection can provide significant insight on the understanding of pathogenesis and immune response. Through the systemic study of the immune response and antigenic protein detection, potential candidates for the diagnostics of the infection in the early stage of disease process can be found. The purpose of this study was to analyze the humoral immune response of the caprine model that was aerosol-challenged with *B. pseudomallei*, mimicking inhalational melioidosis in humans. Melioidosis is common in goats living in melioidosis-endemic locations and the disease in goats has many parallels to human melioidosis [28]. In this present study, we analyzed antibody reactive proteins and determined quantitative humoral antibody response to melioidosis as measured by western blotting and whole cell lysate ELISA. Antibody generation by the humoral response is a key component of understanding the immune response and a foundation of the potential biomarkers and vaccine development. Understanding the progression of immune response through an animal model could give insight into how the host reacts to *B. pseudomallei* infection and what antigens contribute to immune reactivity.

## Methods

### Broad characterization of humoral response

Whole cell lysate (WCL) generated from the *B. pseudomallei* infection strain (MSHR511) was used as the antigenic material to characterize the humoral response in goat sera for 1, 4, and 5 days before infection (pre-challenge) and for days 7, 14 and 21 after infection (post-challenge).

### Design of *B. pseudomallei* aerosol infection (challenge) study and collection of sera

*B. pseudomallei* (MSHR511) isolated from an outbreak of melioidosis in goats on a farm outside Darwin, Northern Territory, Australian goat farm was grown in Muller-Hinton (MH) broth as described in Soffler *et al.* [29]. The bacterium was harvested in mid-log phase and diluted to 1 × 10^4^ CFU/ml as final concentration. The bacteria suspension was then delivered as an intratracheal aerosol [29]. Twelve goats (7 males and 5 females) obtained through a private sale were acclimatized for 1 week before being infected with *B. pseudomallei* under anesthesia (Table 1). The goats were monitored by rectal temperature and complete blood counts pre- and post-infection [29]. Pre-infection sera were available from 8 goats. At the different time points of day 7, 14 and 21 post infection, 2-3 goats were euthanized and sera were collected with the exception of goat no. 16 planned for day 21 but became moribund. The serum from the goat euthanized on day 16 was included with the day 14 for calculations and analysis.

**Table 1.**
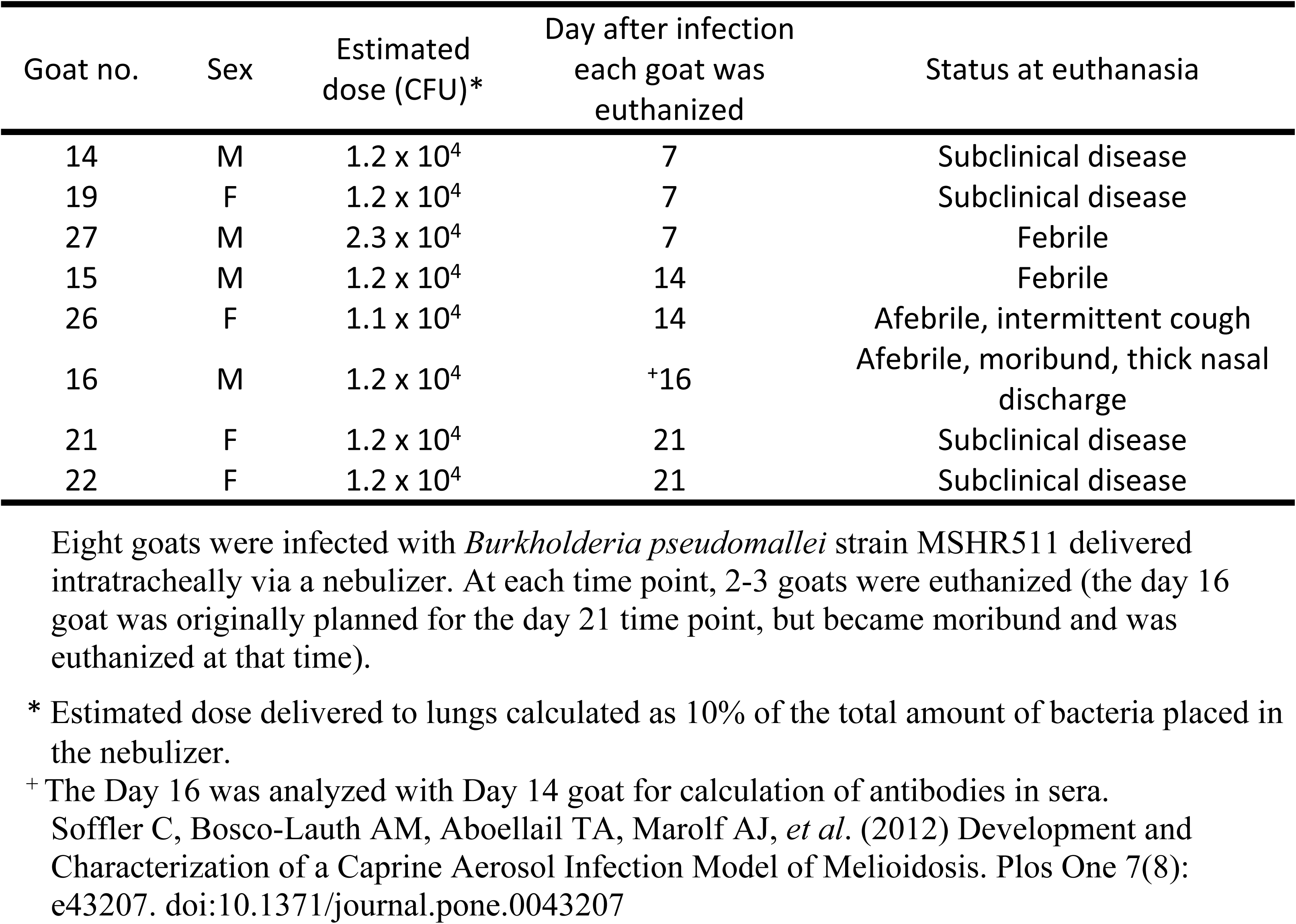
*B. pseudomallei* goat aerosol challenge study design (Soffler *et al*. 2012)

http://www.plosone.org/article/info:doi/10.1371/journal.pone.0043207

### Bacterial Strain and Growth Conditions

*pseudomallei* strain MSHR511 was used in this study. The bacterial strain was grown on minimal media (BD) supplemented with casamino acids and glucose agar plates at 37°C for 35– 48 hr. After incubation, single colonies of bacteria were scrapped and suspended in phosphate buffered saline (PBS) solution, pH 7.4 to give a turbidity reading of 1.0-1.2 at OD 600 nm.

### Purification of Whole Cell Lysate (WCL) Proteins and Antigenic Carbohydrates

WCL proteins were surveyed for immunogenic reactivity using sera from *B. pseudomallei* infected goats by 2-DE western blots. The bacterial cells suspended in PBS buffer, pH 7.4 were washed and centrifuged twice at 16,000 xg for 3 min at 4°C to pellet the cells. The cell pellets were resuspended in lysis buffer (50 mM KH_2_PO_4_, 400 mM NaCl, 100 mM KCl, 0.5% Triton X-100, and 10 mM imidazole), pH 7.4. The bacterial cells were lysed by a freeze and thaw technique using liquid nitrogen and 42°C heat block, respectively, repeated three times. Whole cell lysate proteins were separated by centrifugation at 18,000 xg for 15 min at 4°C. After separation, lysis buffer was exchanged with Tris-HCl buffer, pH 7.8 by centrifugation using microcentrifuge tubes. Protein concentration was determined using the Bradford technique [30] with bovine serum albumen (BSA) as a standard. WCL protein was enzyme treated to remove nucleic acid and precipitated using 15% trichloroacetic acid (TCA) in acetone and centrifuged at 18,000 xg for 18 min at 4°C to purify the proteins. Protein pellets after purification were dissolved in rehydration buffer containing 7 M urea, 2 M thiourea, 1.3% CHAPS, 30 mM DTT, 0.5% NP40, and 0.25% IPG ZOOM^®^ carrier ampholyte of pH 4-7.

For the CPS purification, broth in 2 L baffled Erlenmeyer flasks was inoculated with *B. pseudomallei* RR2683 and incubated overnight at 37°C with shaking (200 rpm). Cell pellets were obtained by centrifugation and extracted using a modified hot aqueous-phenol procedure [31]. Purified CPS antigens were then obtained essentially as previously described [32].

For the OPS extraction and purification, Intron LPS extraction kit reagents were used (Intron biotechnology, South Korea). The cells were collected from agar plates inoculated with B. pseudomallei strain Bp82 and incubated 36 ∼ 48 hr. at 37°C. Cells were collected from the plates and transferred to PBS buffer. Cell pellets were obtained by centrifugation 1300 x g. The cells were lysed using lysis buffer, chloroform, and purification as per manufactures instructions. The precipitated LPS was pelleted and suspended in water and sterility was confirmed by plating on 5% sheep blood in Tryptic soy agar (Hardy Diagnostics, Santa Maria, CA). The LPS was then further purified using proteinase K and 70% ethanol wash and drying.

### Two-Dimensional Electrophoresis (2-DE)

The antibody-reacted proteins from matched silver stained gels were identified by mass spectrometry. The strongly reactive immunogenic proteins were cloned, expressed and used in ELISA assays. 2-DE analyses were conducted using isoelectric focusing (IEF) of whole cell lysate proteins on immobilized pH gradient (IPG) gel strips (7 cm, pH 4-7 NL) for the first dimension according to Rabilloud [33] and Gorg *et al.* [34]. IPG strips were passively rehydrated with proteins dissolved in 165µl of rehydration buffer solution (Invitrogen, Carlsbad, CA) containing 100µg protein. IEF was focused on an electrophoresis apparatus (Xcell6™, LifeTech, Carlsbad, CA) for a total of 8000V.hr. Focused proteins in the IPG strips were reduced and alkylated for the second dimensional electrophoresis by reduction and alkylation buffers for two consecutive 20 min incubations in 100mM Tris-HCl, pH 8.8 containing 5M urea, 800mM thiourea and 4% SDS, alternately, dithiothreitol (DTT) and iodoacetamide (IAA), each with a concentration of 130mM. The second dimension was separated on 4-20% Tris-Glycine gradient SDS-PAGE gels (Novex gels; Invitrogen, Carlsbad, CA). The gels were run at a constant voltage of 110V for 90 min and visualized using silver staining (Shevchenko *et al*., 1996). The images were captured using the UVP gel documentation system (UVP, Upland, CA). Image analysis was performed using Melanie (GeneBio, Geneva, Switzerland).

### Western Blot Analysis

Proteins on 2-DE gels were transferred onto nitrocellulose membrane (Immun-Blot™, 0.2 µm) using a dry transfer technique (iBlot®, Dry Blotting System; Invitrogen, Carlsbad, CA) for 10 min at a constant current of 25 mA. The membrane was blocked for 1h in PBS containing 1.5% skim milk. Blotted samples were reacted sequentially with goat serum at a dilution of 1:1000 in PBS containing 1.5% skim milk for 1h, washed three times in PBS, followed by goat anti-human IgG/IgM horse radish peroxidase at a dilution of 1:1000. Protein spots were visualized using a chromogenic substrate, 3, 3’-diaminobenzidine (DAB) solution.

### In-Gel Trypsin Digestion

Immunostained protein spots were matched with their corresponding protein spots on silver stained 2-DE gels using Melanie software (Melanie version 7.0.6 software; GeneBio, Geneva, Switzerland). The matched protein spots on silver stained 2-DE gels were excised and destained in 0.02% sodium thiosulfate and 0.5% potassium ferricyanide solution [35]. Gel pieces were washed, dried in 50% acetonitrile, reduced and alkylated in buffer (100 mM Tris-HCl, pH 8.8 with 5 M urea, 0.8 M thiourea and 4% SDS) containing 10 mM DTT followed by 100 mM IAA. Proteins were digested overnight in a digestion buffer (50 mM NH_4_HCO_3_, containing 1 mM CaCl_2_) and 12.5 ng/ml trypsin (Promega, Madison, WI) at 37°C. The enzyme treated peptides were extracted using 5% formic acid and 50% acetonitrile. The extraction of the digested peptides was facilitated by vortexing followed by sonication each for 30 minutes.

### MALDI-TOF Mass Spectrometry

Mass spectrometry analysis was performed using a 4800 Plus Matrix Assisted Laser Desorption Ionization Time of Flight (MALDI-TOF) analyzer (AB Sciex, Toronto, Canada) with a 400Å Anchorchip TM target plate (AB Sciex, Toronto, Canada). Recrystallized α-hydroxycinnamic acid (1 mg/ml) in acetone was diluted 1:2 with ethanol and 1 μl was mixed with 0.5 µl of peptides and crystallized on the target. Spectra were analyzed and proteins were identified in a 4000 series Explorer V3.5.3 and Protein Pilot V4.0 software (AB Sciex, Toronto, Canada). Peptide mass fingerprints were searched against RAST annotated protein database. One missed cleavage per peptide was allowed, and the fragment ion mass tolerance window was set to 100ppm.

### Bioinformatics and Protein Identification

Using MS peptide sequence results, “immunogenic” proteins were identified with the help of several software programs available online. Protein identity was performed using BLAST (www.ncbi.nlm.nih.gov) against our laboratory created database and NCBI database was used for inferred bioinformatics information of the identified proteins. Theoretical molecular weight and pI values were taken from NCBI database and calculated using Compute pI/Mw tool on ExPASy website (http://web.expasy.org/compute_pi/).

### Primer Design and PCR

Primers were designed based on the 5’ and 3’ ends of the gene sequences using online software Integrated DNA Technology. Oligonucleotides were generally between 18-24 bases (S1 Table) with a melting temperature of between ≥ 54 – 60°C. PCR was by Pfx DNA polymerase (Invitrogen) [36] to produce blunt-ended PCR products suitable for cloning in pcDNA 3.1 (Invitrogen). PCR reactions consisted of 0.3 mM of each dNTP, 1 mM of MgSO_4_, 3x PCR enhancer solution, 0.3 µM of forward and reverse primers and 0.05 ng/µl template DNA using 1.25U/well Pfx DNA polymerase in a final volume of 20 µl. The PCR amplicons were quantified and analyzed by nanodrop spectroscopy and/or gel electrophoresis and working concentrations of DNA made for use in the ligations reactions. The cloned gene inserts were confirmed by Sanger DNA sequencing [37] and transformed into *Escherichia coli* (strain BL21-DE3).

### Preparation of Recombinant Proteins

#### Bacterial cells

*Escherichia coli* cells were harvested and purified according to the methods previously described [38-40]. *E. coli* cells (strain BL21-DE3) containing his-tagged recombinant proteins were grown at 37°C overnight in 50ml Luria broth containing 100 µg/ml ampicillin. The bacteria were grown in an incubator with an orbital shaker at a speed of ∼200 rpm to log phase (OD_600_ = 0.5 – 1.0). The cells were then induced with isopropyl β-D-1-thiogalactopyranoside (IPTG) of 1mM final concentration.

### Harvesting of *Escherichia coli*

The cells were harvested as a pellet by centrifugation at 4,000 rpm for 40 min at 4°C. The supernatant was discarded and the pellets placed in a dry ice/ethanol bath for 5-10 min. The frozen pellets were transferred into 200 ml of neutral (N)-lysis buffer (50 mM Tris-HCl, pH 7.5, containing 0.3 M NaCl and 0.5 mM EDTA) in a 500 ml flask. Lysozyme (0.5 mg/ml) was added to the solution, incubated for 2 hr to digest the bacteria cell wall. The bacteria cells were then lysed by adding 20 ml of 100 mM MgCl_2_/10 mM MnCl_2_ salt solution. DNA was removed by adding DNase I (10 μg/ml) to digest DNA for 30 min at RT. The resulting bacteria lysate (∼220 ml) was dialyzed overnight with two changes of 4 liter phosphate buffer (20 mM Na_2_HPO4.2H_2_O, 300 mM NaCl), pH 7.4 containing 5 mM imidazole.

### Purification of Recombinant Proteins by Liquid Chromatography

After overnight dialysis, the supernatant was separated from cell debris by centrifugation at 10°C at 10,000 rpm for 30 min. The recombinant proteins were collected as soluble protein. The soluble protein fraction was separated by fast protein liquid chromatography system (BioLogic, Bio-Rad Laboratories, Hercules, CA) using nickel affinity chromatography. The dialyzed protein solution was applied to a 20 ml nickel nitrilotriacetic acid (Ni-NTA) column and eluted with dialysis buffer containing 500 mM imidazole. The purification of His-tagged recombinant proteins was based on the His-tag protocol[41]. The purification of His-tagged recombinant proteins was monitored by SDS-PAGE and confirmed by western blotting using HisG monoclonal antibody, mouse HRP conjugate (Invitrogen). Protein concentration was estimated by Pierce BCA method with BSA as the standard (ThermoFisher Scientific, Grand Island, NY).

### Enzyme-Linked Immunosorbent Assay (ELISA)

To understand the humoral response to *B. pseudomallei* in an experimental goat model for melioidosis, we developed the following ELISA assays to quantify the antibody concentration to the whole cell lysate and each individual protein and cell wall antigens. Five of these were protein antigens (dihydrolipoamide dehydrogenase of pyruvate dehydrogenase complex, PDHD; thiol peroxidase, TPX; alkyl hydroperoxide reductase subunit C-like protein, AhpC2; enolase, Eno; heat shock protein 60 family chaperone, GroEL1) and two polysaccharides (capsular polysaccharide, CPS and type A O-polysaccharide, OPS A) known to be immunogenic in *B. pseudomallei* (S2 Table) [42]. The polysaccharides were selected based on the fact that *B. pseudomallei* produces both CPS and OPS A which are implicated in *B. pseudomallei* virulence [6, 43, 44].

The 96-well immuno plates (Microfluor 2; Fisher Scientific, Pittsburgh, PA) were first coated with individual antigens, namely, recombinant proteins (250ng/well), OPS A (2000 ng/well), or CPS (125 ng/well) in PBS coating buffer (137 mM NaCl, 2.7 mM KCl, 10 mM Na_2_HPO_4_, 2 mM KH_2_PO_4_, pH 7.2) (Fisher Scientific, catalog no. BP3991) and left overnight at 4°C. After overnight coating, wells were washed 4 times with 200 µl PBS. The 96 well plates were flicked between the washes to remove all the PBS from the wells and for the fourth wash the plates were smacked onto a stack of paper towels. After washing, the wells were blocked with blocking buffer solution containing 1% (v/v) BSA in PBS for 2 hr at room temperature (RT). After 2 hr incubation the wells were washed 4 times with PBS containing 0.05% (v/v) Tween-20 (PBS/T) and three different goat sera dilutions in blocking solution added to the wells in a volume of 100 µl per well. Following the 2 hr incubation with the primary antibody (sera from *B. pseudomallei* infected goats), the wells were again washed 4 times with PBS/T to remove unbound goat serum antibody. A secondary antibody of donkey anti-goat IgG or IgM conjugated with horseradish peroxidase (Santa Cruz Biotechnology, Dallas, TX) was added for immunoglobulin G or M detection. Wells were washed 4 times with PBS/T and the enzyme reaction was detected by adding 100 µl substrate (Amplex Red reagent, Life Tech) for a defined time at RT. The fluorescence of the wells was read using a BioTek plate reader at 530/25 excitation and 590/35 emission wavelengths.

## Results

### 1. Initial characterization of total IgG and IgM antibody responses

We initially set out to evaluate the general antibody response to *B. pseudomallei* during an aerosol infection of goats. To do this we determined the relative number of the immune reactive antigenic protein spots as an indicator of the diversity of the antigens the immune response generated to and the quantity of the reactive antibodies. This involved ELISA, 2D gel electrophoresis, and 2D western blot analyses using a whole cell lysate of the infecting strain (MSHR511). Both ELISA and western blot results indicated a much greater humoral response in both breadth and quantity of IgG antibodies compared to that of IgM antibodies throughout the course of infection (Fig 1). Goat IgG antibodies assayed using ELISA on WCL from *B. pseudomallei* MSHR511 showed a rapid increase, rising from an average concentration of 11,348 µg/ml on day 7 to 26,666 µg/ml on day 14 and reaching a maximum of 35,088 µg/ml by day 21. The concentration of IgM goat antibodies increased over time as well, from a mean of 4,026 µg/ml on day 7 to 8,565 µg/ml on day 14 and reaching 9,590 µg/ml by day 21. The total amount of goat humoral IgM antibody was 2-4-fold less than goat IgG antibodies for all three-time points after challenge with *B. pseudomallei*. In the western blots, the relative number of IgG-reactive protein spots detected after being probed with goat sera mirrored the increase observed using ELISAs for days 7, 14 and 21 (Fig 1). However, the number of antigenic WCL protein spots on IgM western blots remained relatively the same across all time points, with an average of 18 on Day 7, 19 on Day 14, and 12 antigens on day 21. The difference of the IgM immune response between ELISA and 2D western blot might be explained by the particular antigens used by the two methods to detect antibodies. ELISA used whole cell lysate, which included both proteins and carbohydrates, while the western blot used only the whole cell lysate proteins, although this would include glycoproteins. CPS and OPS are carbohydrate antigens known to trigger strong IgM reaction and would only be detected by the WCL ELISA. Overall, the *B. pseudomallei* aerosol challenge induced an increasing and diverse antibody response over the disease progress in the goats.

**Fig 1.**
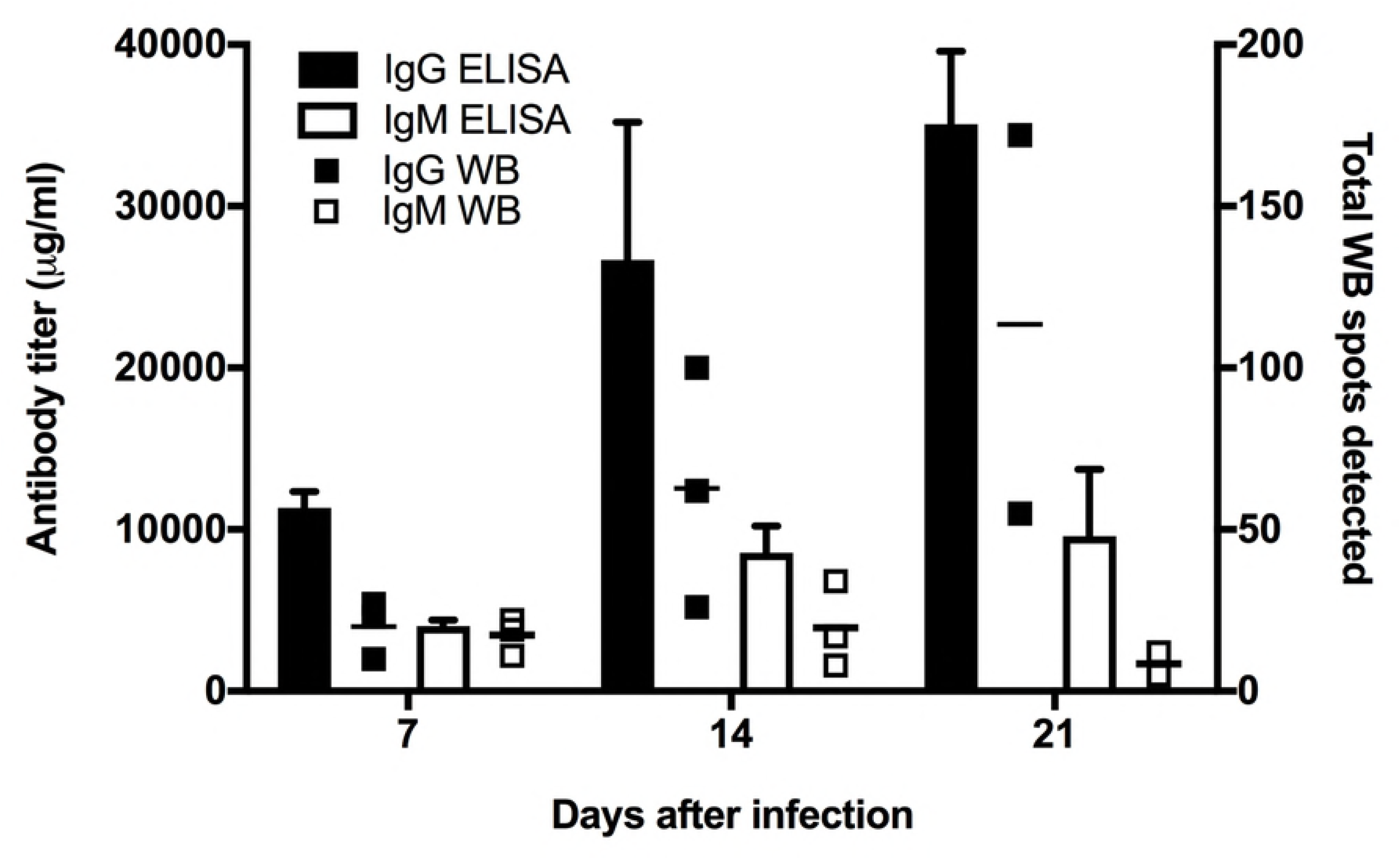
Quantitative goat humoral antibody responses to melioidosis as measured by whole cell lysate (WCL) ELISA and western blots.

Goats were challenged by aerosol infection with Burkholderia pseudomallei isolate MSHR511. ELISA and western blot analyses were performed using whole cell lysate (WCL) of cultured MSHR511. Goat sera were drawn prior to challenge and on days 7, 14 and 21 after infection. ELISA results (left y-axis) show the mean antibody titer from all goats per sampling day (7, 14, or 21) and are represented by vertical bars for IgG ▬, and IgM ▭. The amount of goat antibody in micrograms (µg) was calculated by subtracting pre-challenge from post-challenge ELISA results (before averaging) relative to the commercially purchased purified standards of goat IgG and IgM. The right y-axis shows the total number of antigenic spots in western blots from each goat, represented by small squares for IgG ■, and IgM □. The total numbers of antigenic spots for IgM and IgG are listed in Table 1.

### 2. Broad survey of goat IgG and IgM responses of pre- and post-infection (challenge)

We characterized the IgG and IgM antibody responses of 8 individual goats’ serum samples using 2D electrophoresis and western blots. Our 2DE reference maps prepared from MSHR511 WCL allowed us to detect nearly 600 bacterial protein spots by silver stain and to match those spots with antigenic spots detected by IgG and IgM western blots (Sl Fig), thus providing a large population of proteins to investigate. High numbers of immunogenic protein spots were detected (marked in blue), denoting a highly diverse humoral response to bacterial proteins. Immunogenic protein spots detected on the western blots showing strong immunoreactivity against IgG increased in intensity over the 21 days of infection. In contrast, immunogenic protein spots for IgM showed low intensity and remained that way for the entire infection period (Fig 2). Even 2D western blots probed with pre-infection sera showed multiple immunogenic protein spots reactive for IgG (0-20 spots) and IgM (0-12) in most goats (Figs 3 and 4). This suggests that the humoral antibody response was previously primed to recognize these or similar antigens from prior bacterial exposure. After infecting the goats with aerosolized *B. pseudomallei,* the number of immunogenic protein spots reacting with goat IgG showed a marked increase over day 7, 14, and 21. The average number of IgG spots shows an increase from 29 protein spots at day 7 to 74 on day 14 and 129 protein spots on day 21 (Fig 3).

**Fig 2.**
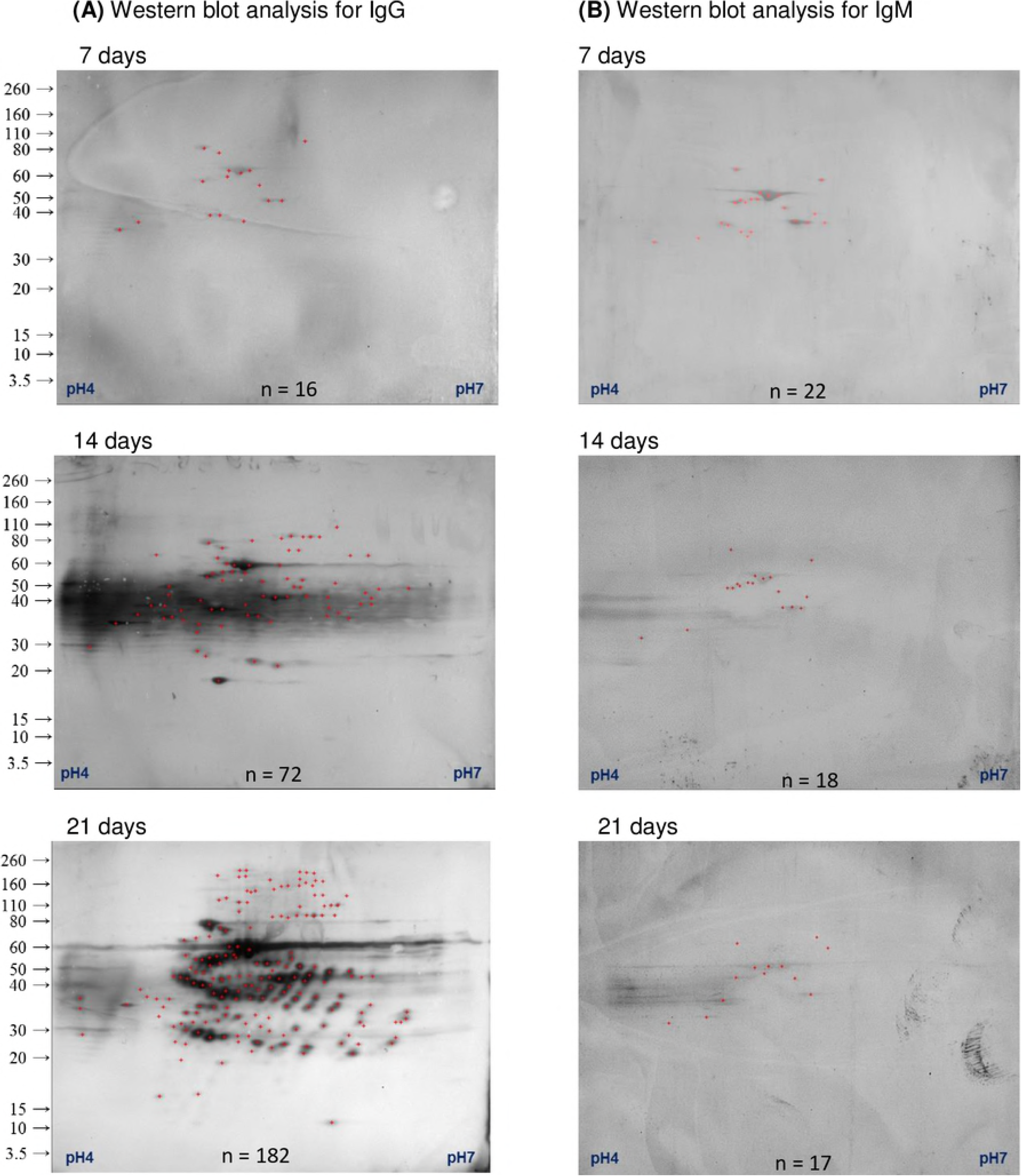
A time series of IgG and IgM responses to infection with *B. pseudomallei*.

Goat sera were drawn prior to infection and on the day of euthanasia. Immunoreactive proteins with IgG (A) and IgM (B) were determined by western blot analysis and then mapped onto a silver stained gel. The number of immunogenic protein spots detected (n) is provided at the bottom of each image.

**Fig 3.**
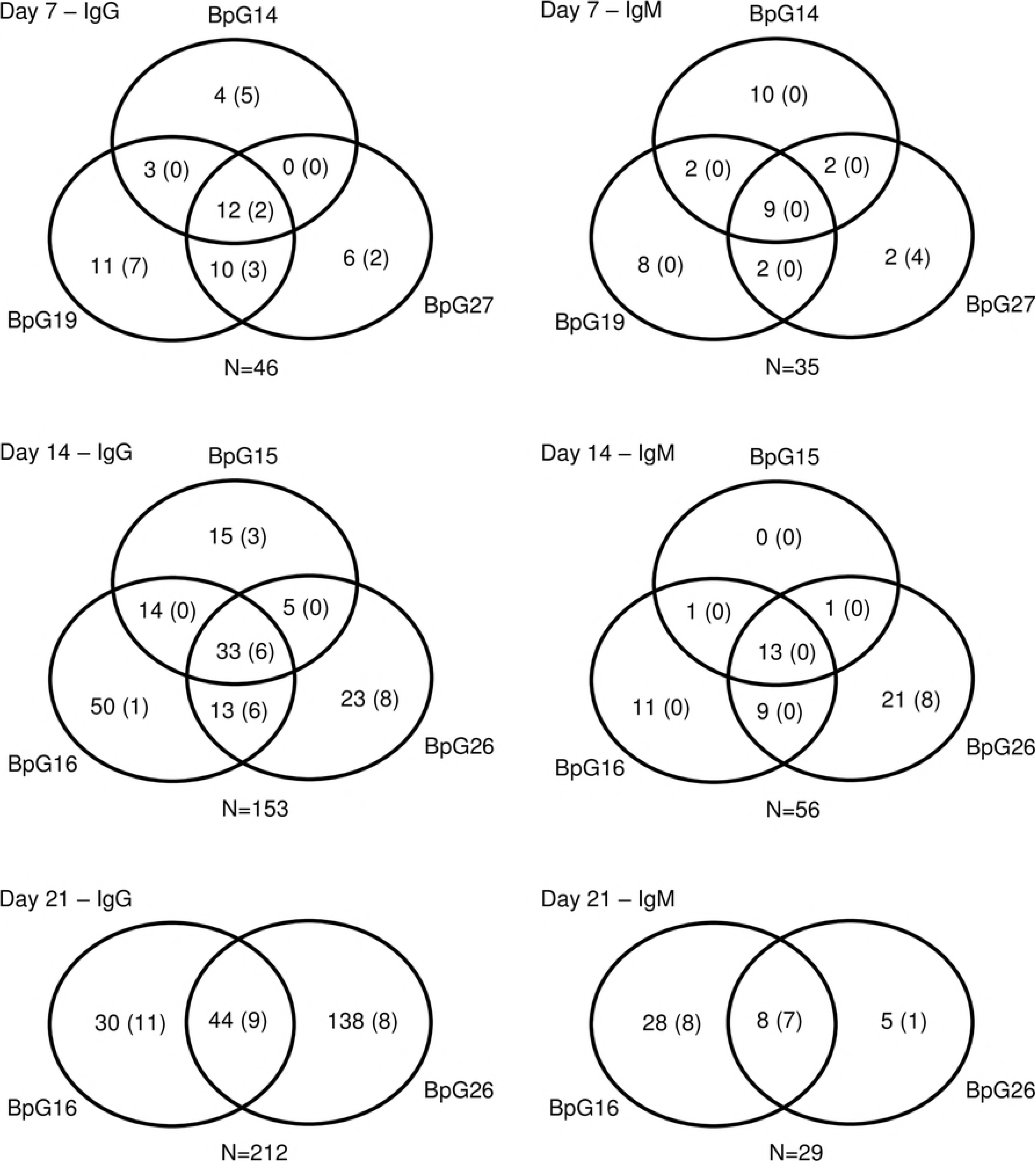
Comparison of constant and variable immunogenic proteins among individual goats.

All the immunogenic protein spots were identified using Burkholderia pseudomallei MSHR511 whole cell lysate protein and eight different goat sera collected during the time course of infection; day 7, 14, and 21 after challenge. All the detected antigenic protein spot count is 282. The number inside each circle denotes the total number of unique detected immunogenic proteins at a particular time point; numbers in parentheses are the total number of pre-challenge immunogenic proteins for that particular goat. The total number of immunogenic proteins detected for each isotype and time point is indicated (N)

**Fig 4.**
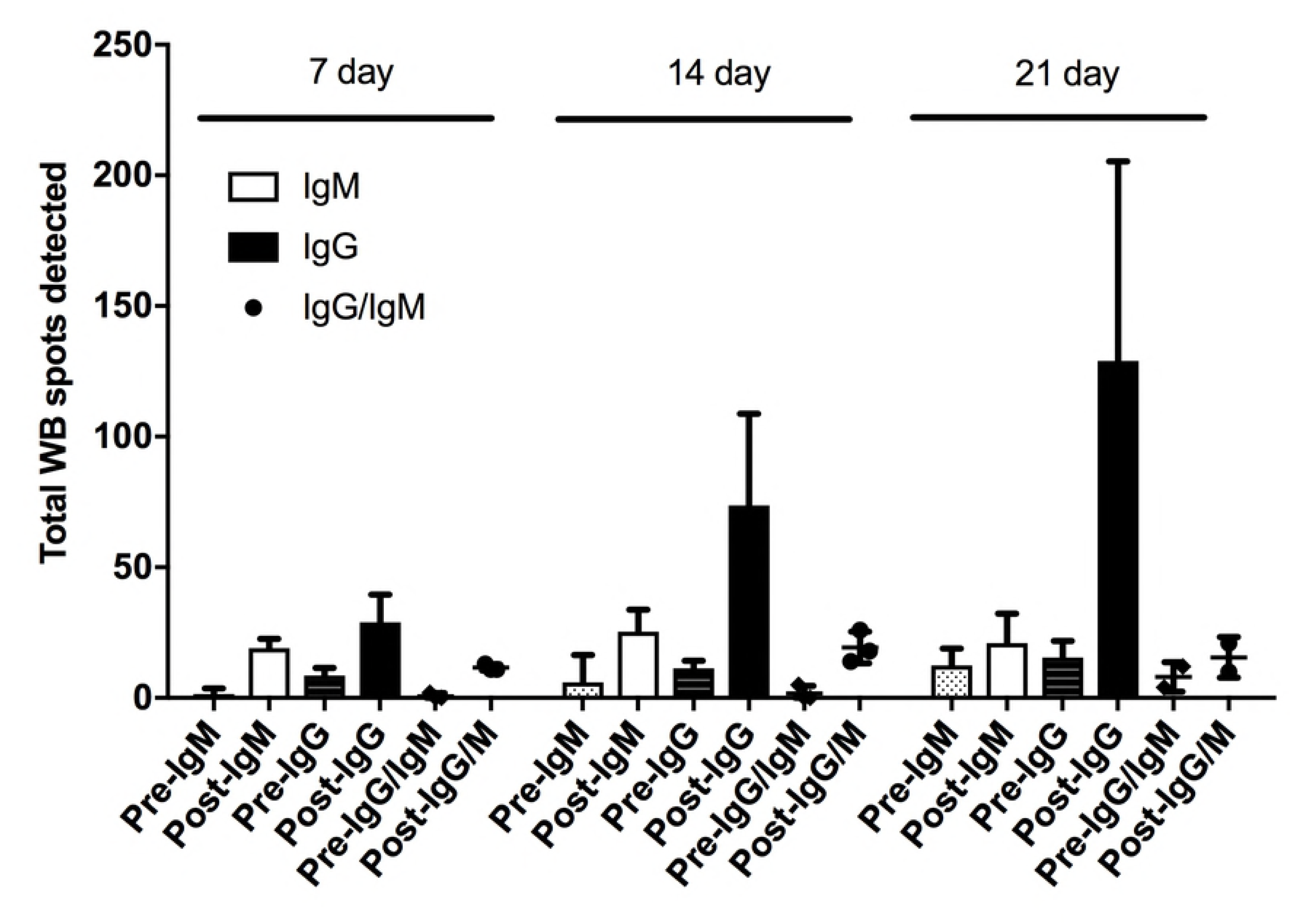
Comparison of IgG and IgM Immune diversity across eight infected goats against proteins of Burkholderia pseudomallei MSHR511.

The average of antigenic spots detected by each individual antibody isotype is shown for each collection time after aerosol challenge and the paired pre-challenge serum sample; 7, 14, and 21 days. In addition, spots that were detected by IgG and IgM antibodies were averaged with pre- and post-challenge immune diversity. Vertical Bars represent immunoglobulins M (IgM) ▭, and immunoglobulin G (IgG) ▬. Circle represents spots detected by both of IgM and IgG ●.

The number of IgM immunogenic protein spots was much lower compared to IgG. At the pre-infection time point, a mean of about 6 IgM antigenic protein spots were detected but all were very faint (Fig 2). The number of IgM immunogenic protein spots increased 3-fold by day 7 (mean of 19 spots), but then remained stable for most goats with a mean of ∼25 IgM immunogenic protein spots on day 14 and 21 immunogenic protein spots on day 21 (Fig 4). This result was in agreement with that total IgM response to MSHR511 WCL (above) grew weaker over the 21day time interval.

### 3. Comparison of constant and variable immunogenic proteins

Over the time course of this challenge study, we found that a subset of proteins raised a consistently strong immunogenic response at each time point for all individual goats, especially for the IgM response: heat shock 60 family protein GroEL, elongation factor -Tu, ATP synthase beta chain, and heat shock protein DnaK. Many antigenic proteins were observed to remain reactive after the initial response was seen. However, there were also many antigenic proteins observed at only a single time point per goat. Considering all of the antigens detected for IgG and IgM, there were 44 antigenic proteins found at all-time points post challenge, 14 of which were also observed in pre-infection sera (Fig 3). There were 6 protein spots that showed reactivity to both of IgG and IgM. Of all of the antigens for IgG, 38 protein spots were detected at all of the post-infection time points, making proteins reactive to IgG antibody as the most common (38 out of 44 proteins). Eight of these antigens were found in the pre-infection sera (Fig 3 and S Table 4). On the other hand, IgM had the least number of antigenic proteins spots present at all-time points with 19 out of 44 proteins and only 5 antigenic proteins reactive in the pre-infection sera (Fig 3). The total WCL reactive IgM was stable with the majority of antigens (19) present throughout Day 7 -14.

To place our results in a broader context, we compared the goat immune response in this study with antigens previously described in human melioidosis patients [42]. *B. pseudomallei* infection leads to strong immune responses in both goats and humans, and many antigens elicited a comparable antibody response. According to these data, there are 98 antigenic proteins detected in both of human and goat immune responses, which is more than half the number of antigenic proteins in the human immune response. Considering the total amount of antigens detected with goat IgG and IgM, 98 out of 282 is still higher number in comparison with 135 total detected human antigenic proteins (Fig 5). The difference in the number of the detected antigenic proteins might be caused by the various biological facets including anatomy, pathophysiology, and genetics, resulting in the different disease progress due to dissimilar route, dose, and immune state prior to infection in/between human infection and animal model. So, the highly common antigenic proteins detected in both human and animal melioidosis imply that this goat model study could be an appropriate animal model to understand disease progress and humoral immune response with mimicking human disease condition.

**Figure 5.**
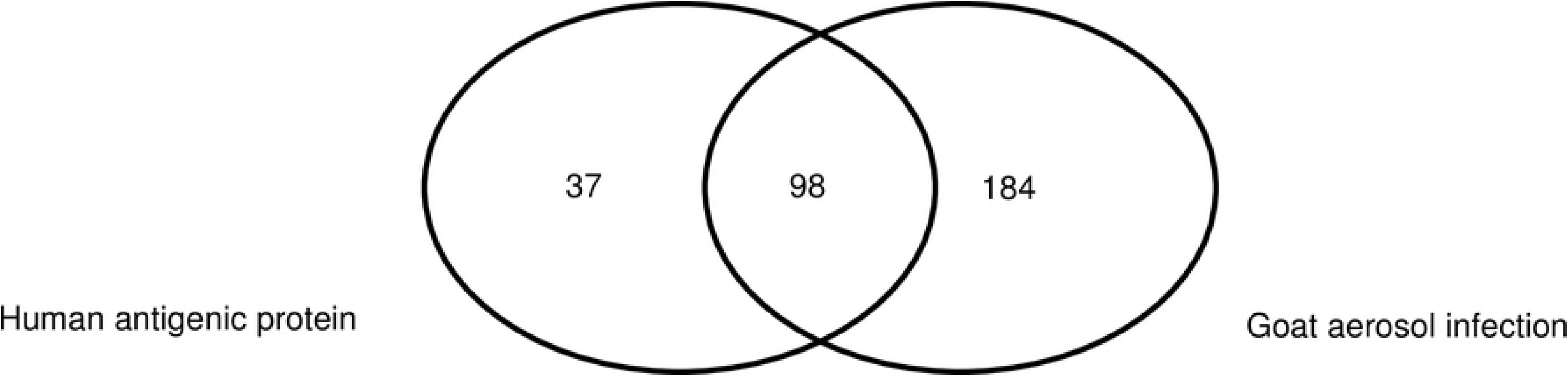
Comparison of immunogenic protein spots common with and specific to goat infection sera and human melioidosis patient sera.

Immunogenic whole cell lysate proteins from B. pseudomallei MSHR511 were detected by western blot and sera from human sera (note: human protein list was taken from 4 melioidosis case paper [42]). The number inside each circle denotes the total number of detected immunogenic protein spots from human and goat sera. Each number denotes human specific immunogenic proteins, common proteins detected both of human and goat sera, and goat sera specific immunogenic proteins at all the time points after the infection.

### 4. Characterization of pre-infection immunogenic proteins

Antibodies to a subset of identified *B. pseudomallei* antigens were found in pre-infection goat sera. We compared these 17 antigens from MSHR511 against a subset of common Gram-negative bacteria that could possibly be the source of a pre-challenge infection, including *Pseudomonas aeruginosa, Escherichia coli, Campylobacter jejuni*, and *Mycobacterium tuberculosis* (Table 2). None of these 17 *B. pseudomallei* MSHR511 proteins had an amino acid identity above 83% when compared to their homologues in the selected species of Gram-negative bacteria. Amino acid identity was highest between MSHR511 and *P. aeruginosa*, with 13 antigens in the range of 55-83% and four with much lower similarity (33-40%). Eight proteins from all of the four bacteria evaluated had a high identity of > 50% compared to *B. pseudomallei*: ATP-dependent chaperone protein (ClpB), elongation factor G2 (EFG2), chaperone protein (DnaK), heat shock protein 60 family chaperone (GroEL1), ATP synthase alpha chain (Atp), ATP synthase beta chain (AtpD1), Translation elongation factor Tu (EF-Tu), and enolase (Eno) (Table 3). The remaining 9 proteins out of 17 were the least similar to *B. pseudomallei* proteins across the four Gram-negative bacteria, with one exception, viz. *C. jejuni*, S-adenosylmethionine synthetase having < 50% amino acid sequence identity to the *B. pseudomallei* protein. *C. jejuni* had the lowest number of proteins with similarity to *B. pseudomallei*, with only eight proteins out of 17 having > 50% amino acid sequence identity to *B. pseudomallei* proteins. Whether the < 50% amino acid identity of *B. pseudomallei* to *jejuni* means reduced immune reaction to the *C. jejuni* protein antigens and its protein epitopes is unknown. Interestingly, amino acid similarity for GroEL was moderately high across all species (58-73%) and may provide insight into the source of the pre-challenge antibody responses we observed.

We compared antigenic spots that were detected using pre-infection sera to the immune reactivity over time. We found that antigenic protein spots detected before the aerosol challenge were also detected at time points after the *B. pseudomallei* aerosol challenge. The chaperone and cell division related proteins in particular were detected pre- and post-infection with high signal intensity (S4 Table).

We chose to more fully investigate the antibody response to six proteins that induced an antibody response prior to *B. pseudomallei* infection. Using 2D western analysis and spot area (a relative measure of immune reactive intensity), each antibody response appeared to peak at a specific time point after the aerosol infection. Goat sera probed against the IgG western blots showed that the antigenic spot areas for each protein antigen (GroEL, EF-Tu, AtpD1, DnaK, AtoC, and Eno) generally increased through day 21 (Supplemental Figure S2). This paralleled the pattern observed for the total IgG response to WCL (above). In addition, the antibody response to these antigens occurred by seven days after infection. In contrast, IgM antibody responses for these six antigens typically showed an early peak in spot area on day 7, which then decreased slowly throughout the remainder of the challenge study. The day 7 peak was particularly strong for GroEL and EF-Tu, and both of GroEL and EF-Tu, and AtoC induced a magnitude of response on par with the IgG antibody at that time point. Interestingly, none of the IgM response for these six antigens matched the overall pattern observed in the total IgM count (i.e., none increased from day 7 to day 14). This pattern indicates that the IgM response to other antigenic proteins must generally grow stronger over infection and may represent new antibody responses.

**Table 2.**
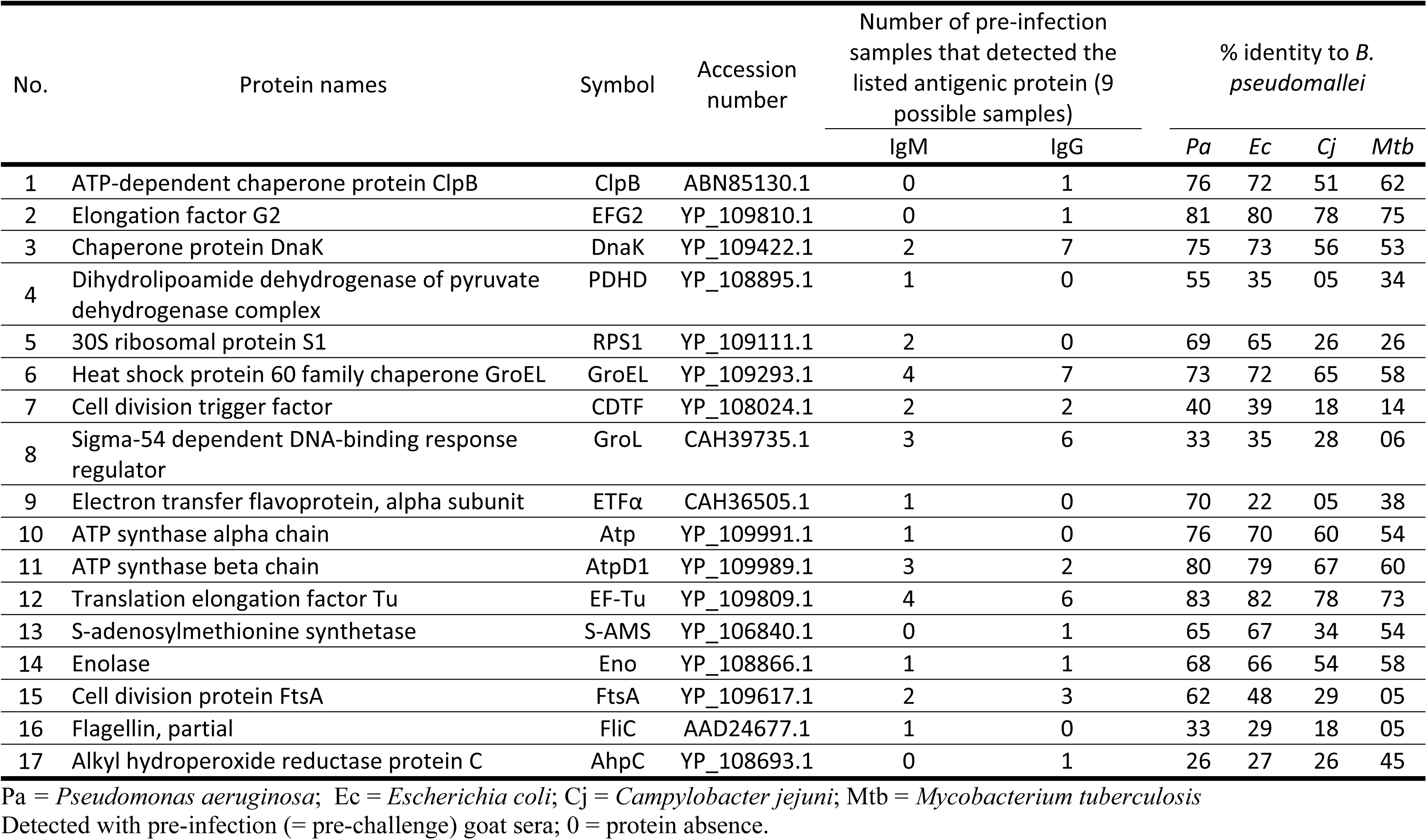
List of identified antigenic proteins detected in pre-infection sera and their percent identity to four other Gram-negative bacteria.

### 5. Using seven highly immunogenic antigens to quantify immune responses

We used ELISA assays to more thoroughly investigate seven antigens (PDHD, TPX, AhpC2, Eno, GroEL, CPS and OPS A) that were immunogenic in this goat study and a previous human melioidosis study including 4 patients (Fig 6). The overall reactivity against IgG increased over the time. The amount of IgG antibody detected on day 14 for 4 antigens (PDHD, AhpC2, Eno and CPS) ranged from 110.8 to 133.2 µg/ml, with the remainder antigens having < 48.9 µg/ml (Fig 6). The exception was GroEL. IgG response specific to GroEL were at a concentration of 329.8 µg/ml on day 7 and reached up to 5579.1 µg/ml by day 21 (Fig 6). The concentration of IgM on the other hand, increased mostly from day 14 for CPS with 28.4 µg/ml to 65.2 µg/ml for GroEL antigen. The 3 antigens showing high levels of antigen-specific IgG antibodies compared to IgM by day 21 were AhpC2, Eno and GroEL. In contrast, the antigens; PDHD, TPX, CPS and OPS A on day 21 had elicited fairly high concentrations of IgM-specific antibodies compared to that of IgG antibodies (Fig 6). The percentage of individual antigens for the goat antibody immune response was calculated relative to the total immune response of the 7 selected antigens (Supplemental Figure S3). IgM antibody responses constituted the highest percent of the immune response with the exception of GroEL antibody immune response, which had very similar percentages for both IgG and IgM antibodies for days 7 and 14 and a significantly high level of IgG goat antibody response for day 21 (Supplemental Figure S3). Because GroEL showed the highest immune response for both IgG and IgM antibodies, we compared the individual antigens antibodies immune response to GroEL (Supplemental Figure S3). The heat shock protein GroEL elicited the strongest goat antibody immune response compared to the other six antigens (PDHD, TPX, AhpC2, Eno, CPS and OPS A) measured in this study, specifically for IgG. This response may be due to prior exposure of the goats to GroEL protein found in other Gram-negative bacteria which may possess conserved amino acid sequences similar to *B. pseudomallei* GroEL. For goat IgM antibody immune response, three antigens, CPS, GroEL and OPS A showed the strongest immune responses for day 7. The other antigens (PDHD, TPX, AhpC2 and Eno) produced similar goat IgM antibody responses for day 7. However, for day 14 the proportion of goat IgM antibody immune response was very similar for all the 7 antigens (Supplemental Figure S3). But by day 21, CPS showed the highest goat IgM antibody response compared to the other antigens; GroEL1, PDHD, TPX, AhpC2, Eno and OPS A. There was not much change in IgM antibody response for both GroEL and OPS A from day 14 to day 21; while the antibodies to the other antigens remained at similar levels for day 14 and 21.

**Figure 6.**
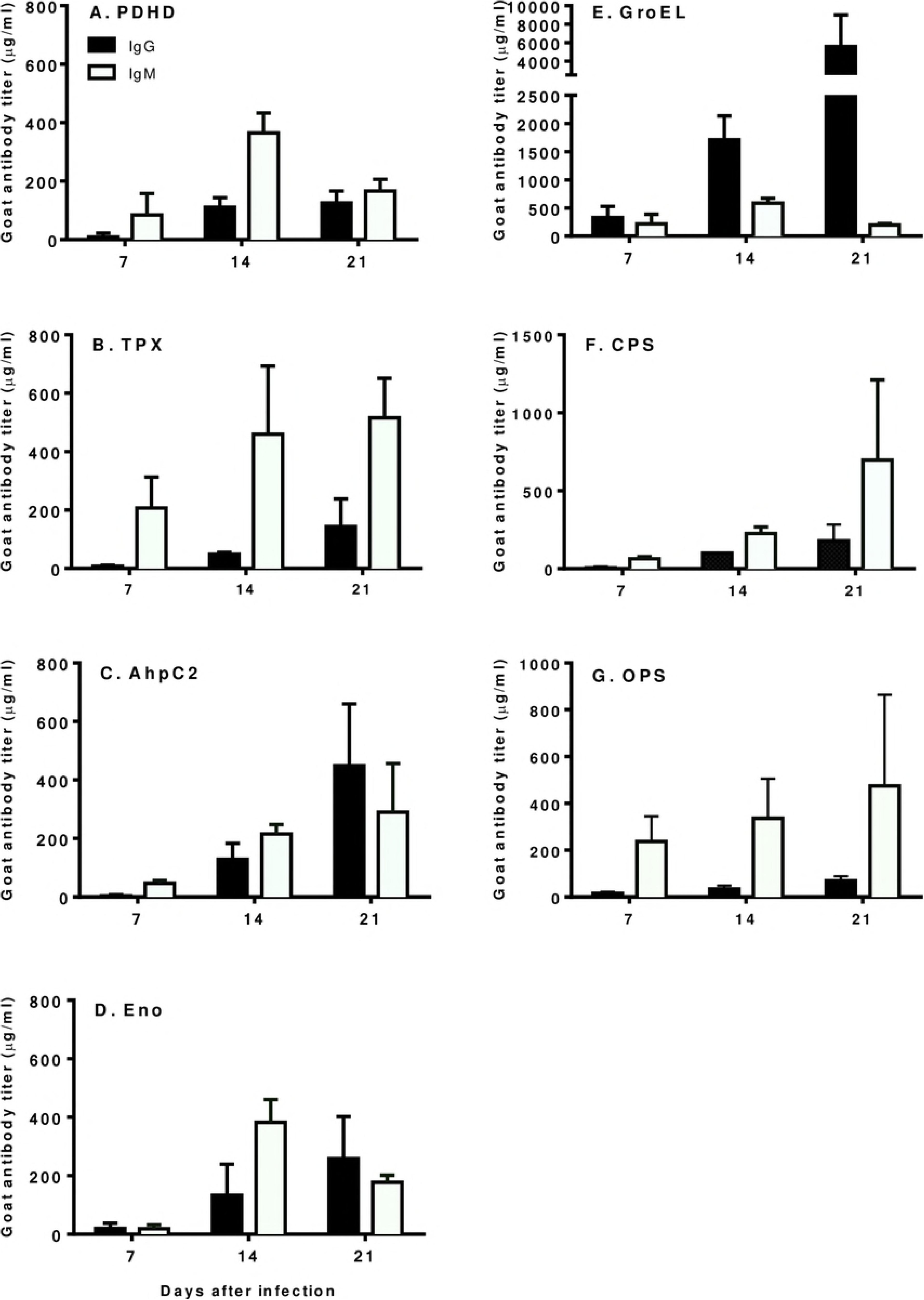
Goat humoral antibody responses to individual protein and polysaccharide antigens.

Goat sera were drawn prior to challenge and on days 7, 14 and 21 after infection (goat serum from day 16 was averaged with the day 14 calculations). Goat antibody quantities (µg/ml) was calculated by subtracting pre-challenge from post-challenge ELISA results relative to the commercially purchased purified standards of goat IgG and IgM. (A) PDHD, Dihydrolipoamide dehydrogenase of pyruvate dehydrogenase complex; (B) TPX, Thiol peroxidase; (C) AhpC2, Alkyl hydroperoxide reductase subunit C-like protein; (D) Eno, Enolase; (E) GroEL, Heat shock protein 60 family chaperone; (F) CPS, Capsular polysaccharide; and (G) OPS A, Type A O-polysaccharide.

## Discussion

### Overall immune response to the challenge

The overall humoral immune response of goats showed an increase of antibody intensiy and immune diversity during the infection progression, especially for IgG. In contrast, IgM reaction showed only a slight increase in antibody intensity and the immune diversity decreased after 14 days post challenge. The overall immune response intensity increase of IgG from the earliest stages of infection implies that there are pre-existing immune conditions for many of the antigens including individual proteins tested in this research. The IgG immune response was greater than the intensity for IgM reactivity. IgM is the primary adaptive immune response to infection while IgG usually develops in the later stage of the infection course after class switching from IgM. However, our investigation showed that the overall IgG humoral response was stronger than IgM at day 7 (initial assay date after goat aerosol infection). This stronger intensity of the IgG response may be explained by protein epitope ability to induce a strong IgG antibody response to protein epitopes of pre-existing conditions such as memory B-cells or pre-primed antibodies which could be produced by cross-reactive epitopes [45].

The immune diversity results as determined by western blot analysis showed a slightly different pattern from the overall humoral response determined by ELISA. The count of IgM immunoreactive proteins showed a similarity to IgG reactive protein spots on day 7 and decreased with the decreased intensity of IgM reactive protein spots over the infection progress (Figure 2 and Figure 2S). At day 14 onward, we detected more IgG reactive proteins than IgM reactive ones (Figure 1 and 2). The IgG antibody demonstrated more unique immunoblot protein spots from day 14 to 21, whereas, IgM reacted with more protein spots on day 7 compared to IgG (Figure 3).

The antibody isotype switch happens between day 7 and day 14 when an antigen(s) persists [46, 47]. Even though there are more detected IgM reactive protein spots on day 7, the intensity of humoral immune response was stronger to IgG. The strong IgG response at the early infection stage like day 7 may arise from the memory cells and prior existing health conditions, which produce faster and stronger humoral immune response with the current infection. The immunogenic proteins sequence comparisons of the selected bacteria showed low similarity to the selected immunogenic proteins of the infecting strains, which could produce memory cells for the fast and strong immune response. However, there is the potential that conserved sequence or structure epitopes of the selected immunogenic proteins between the selected bacteria and the infecting strain or other antigens cause fast and strong IgG response among the unexamined antigens. The T helper cells contribute the class switch and affinity maturation for stronger immunoreactivity. Other researchers have reported similar results of elevated IgG and IgM post infection with *B. mallei*, a closely related species to *B. pseudomallei* [48].

### Immunity of commonly detected protein

We found 44 antigenic proteins commonly detected in all of the individual goats, and 8 immunogenic proteins detected in pre-infection sera for IgG and IgM using western blot. Those immunogenic proteins are a small portion out of the total 282 immunogenic proteins identified in this research (S4 Table). Each or several goat specific immunogenic proteins were detected many more than the commonly detected immunogenic proteins. These results demonstrate how variable individual immune responses are to even infection of the same infecting strain with the same conditions. The individual animals may vary in their prior exposures to other bacterial infections. In addition, these are outbred animals with different genetics, perhaps creating different immune responses because of the polymorphic differences of MHC alleles. However, commonly detected immunogenic proteins with the different timelines for antibody characterization generate the antibody response to the same or similar epitopes despite the host variability to infection. These common immunogenic proteins are mainly immunodominant proteins (S4 Table) [49]. These common immunogenic proteins give the insight of how the host generally reacts to the infection and it also demonstrates those proteins’ potential as general diagnostic antigens for bacterial infection.

### Highly immunoreactive antigens

By investigating and identifying highly immunogenic proteins, we found that five proteins (PDHD, TPX, GroEL, AhpC2, and Eno) and two polysaccharides (CPS, OPS A) that showed high immunoreactivity to melioidosis patient sera against four *B. pseudomallei* strains in our previous study (Yi *et al.*, 2016). Those highly immunogenic proteins were also commonly detected proteins using sera from the eight different goat individuals. The western blot and ELISA results for the selected antigens showed a strong ELISA signal for IgG and IgM antibodies from day 14 onwards. Specifically, as for the proteins, GroEL, AhpC2, and Eno induced a stronger IgG response, while the IgM response was strong for all five antigenic proteins (Figure 6). This is probably due to antigenic protein epitopes inducing a stronger IgG antibody response with class switching after an initial immune response [50].

Polysaccharides, namely CPS and OPS A were assessed in this study. CPS showed a strong ELISA signal for IgG and IgM antibodies from day 14 onwards. While OPS A antigen demonstrated a good IgM antibody response from day 7. CPS and OPS A are thymus-independent antigens known to activate B-cells to elicit low-affinity IgM antibodies [45]. Therefore the presence of IgM up to 21 days in goat sera can likely be explained by the persistence of polysaccharide antigens for an extended period in the lymphoid tissues, continually stimulating newly maturing B-cells to produce IgM antibodies [46].

As is evident from the results, there were differences in the levels of goat antibody titers expressed against the seven highly immunoractive antigens, which may suggest possible differences in the amounts of each protein and polysaccharide antigen produced by *B. pseudomallei* or differences in antigen immunogenicity. Vasu *et al*. [50] has shown that the primary antibody response to *B. pseudomallei* protein and polysaccharide antigens in melioidosis patients was IgG, subclasses IgG1 and IgG2 antibodies, suggesting a Th1 antibody response in both septicemic and non-septicemic melioidosis cases.

As the most immunoreactive antigen, GroEL is one of the immunodominant antigenic proteins, which gave a strong immune intensity even at the day 7 (Figure 6). This result could be explained by the presence of memory B cells from a prior non-melioidosis infection with a related organism and may cause a strong immune response rapidly right after the host encountered the antigenic proteins. Amemiya *et al*. [51] using ELISA assays reported a 10-fold increase of IgG antibodies against the heat shock protein (hsp), GroEL. This is in agreement with our results, where GroEL specific goat antibody titer was ∼12.8-fold higher than it was to AhpC2 protein; the second highest goat IgG antibody titer to GroEL (Figure 6).

Because of this strong IgG and IgM antibody immune response to GroEL, it was decided to examine this result further. First, GroEL family of proteins is reported to contain epitopes that are highly conserved from prokaryotes to humans [52]. GroEL-like proteins are reported to be immunodominant antigens from infectious pathogens, such as *Mycobacterium leprae, M. tuberculosis, Coxiella burnettii* and *Legionella pneumophila* [52]. During infection, a pathogen undergoes selective pressure which increase microbial heat shock protein synthesis to withstand the harsh environment inside the host [53]. This reason is why hsps are major antigens in infectious agents that induce a strong innate and cellular immune response [54].

### Prior immune condition and potential cross-reactivity of selected antigens

Most of the pre-infection sera showed some IgM and IgG immune reactivity to *B. pseudomallei* proteins. Most of the antigenic proteins identified using pre-infection sera were immunodominant proteins [42], and were mostly faint spots of low intensity. Strongly reactive proteins to IgG from the pre-infection sera set were detected, which were mostly immunodominant proteins and also detected at post-infection sera (S4 Table). The goats showing pre-primed immune reactivity showed fewer detected antigenic proteins, thus exhibiting less immune diversity (Figure 4). The results imply that the pre-primed response and immunodominant antigens impede developing a broader antibody response against multiple epitopes of the strain. Those antigens may be cross-reactive with proteins from prior bacteria that the goat came in contact with.

As observed with other diseases, potential serological cross-reactivity to proteins of closely related pathogens can produce the immunological memory of the B-cells and cause cross-reactivity in immunoassay, presenting a diagnostic challenge. This cross-reactivity was investigated by comparing the amino acid sequences of seventeen antigenic proteins with the same proteins found in a selected number of Gram-negative bacteria, viz. *P. aeruginosa, E. coli, C. jejuni* and *M. tuberculosis* (Table 2). The percent identity of *B. pseudomallei* proteins to the above four Gram-negative bacteria ranged from 5% for PDHD and ETFα of *C. jejuni,* FtsA and FliC of *M. tuberculosis* to 83% for EF-Tu of *P. aeruginosa.* The genus, *Pseudomonas* was where *B. pseudomallei* was classified before *Burkholderia* was proposed as a new genus [55, 56]. The overall sequence comparison results showed the low similarity of the selected proteins of the chosen bacteria even though the investigated proteins are commonly detected proteins among the bacteria. The low sequence similarity might indicate that the studied proteins do not contribute to the early stage of immediate antibody response with high humoral response. However, there is potential that there are still cross-reactive epitopes or unstudied antigens causing cross reactive humoral response at the early challenge stage. The faint IgG and IgM immunoreactive protein spots detected in this study support the postulate. Overall, the selected proteins even showed cross reactivity in this research but the intensity and the sequence similarity were very low. Thus, the selected proteins could be the potential biomarkers of *B. pseudomallei* infection (Table 2).

In conclusion, this study characterized the overall antibody response of IgG and IgM antibody response, delineating the diversity of immunogenic proteins generated within the host after *B. pseudomallei* aerosol challenge. There was a detectable immune response from the early stage of the infection and there are antigens eliciting strong signal intensity for either/both of IgG and IgM. Of the antigens detected, there were 44 commonly detected antigens among the eight individual goats (S4 Table). Many of the detected antigens demonstrated the variation of the immune responses among the goats and during the infection progression. This study also involved expression and purification of five recombinant proteins and 2 polysaccharides detected to be immunogenic using sera from *B. pseudomallei* infected goats and also from human patient sera. Of the seven antigens assayed, AhpC2, Eno and GroEL had a stronger IgG response, while CPS, OPS and TPX showed the stronger the IgM response. Even though we tested the seven potential antigens for their immunoreactivity and the potential as diagnostics biomarkers, we detected additional antigenic proteins with potential as diagnostics targets showing high detection frequency among the goats. Further ELISA assay evaluation of these antigens is needed to determine if they are an improvement over the IHA assay sera and in clinical settings.

## Supporting Information Legends

**S1 Fig.**
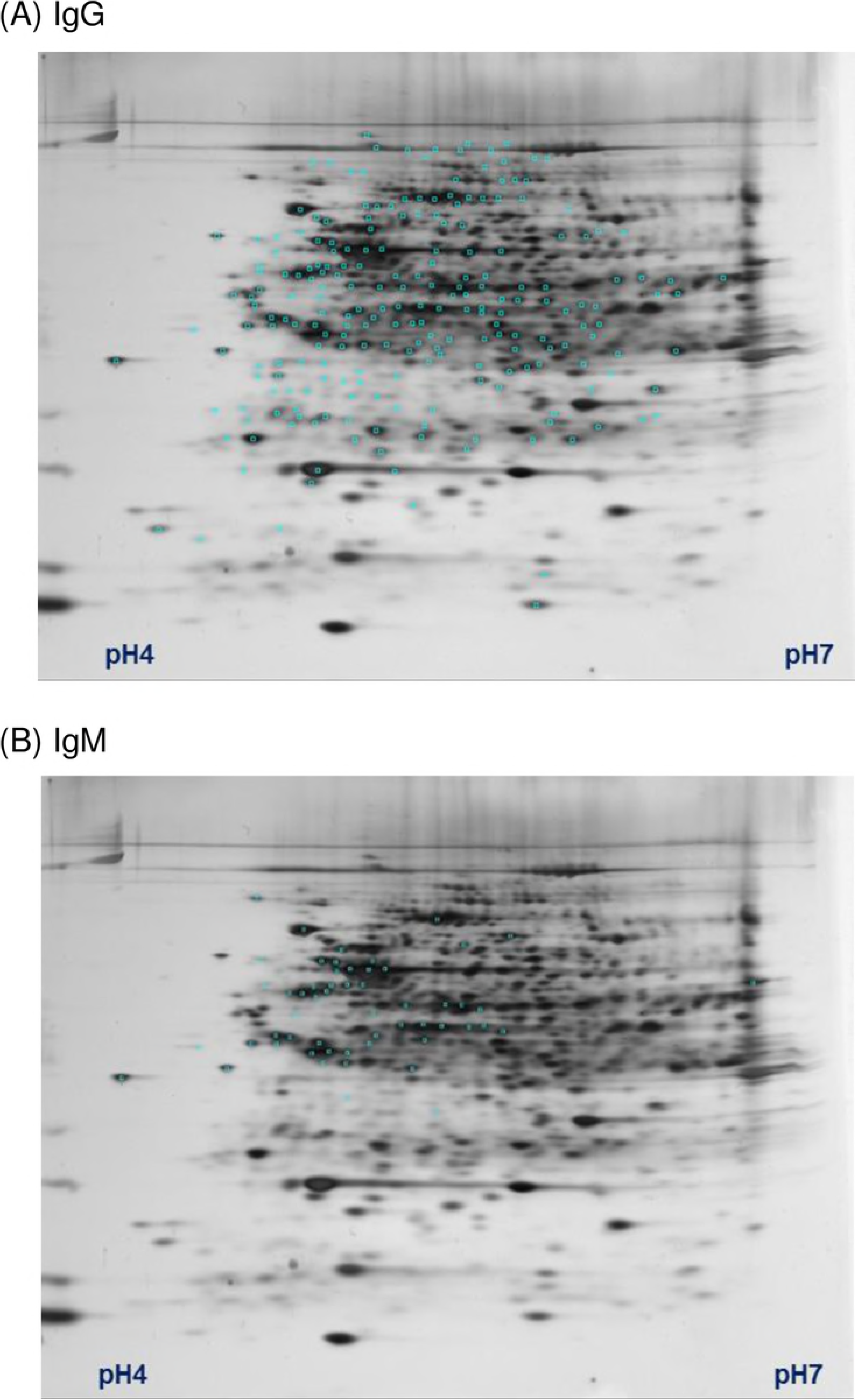
Electrophoretic reference maps of proteins cross reacting to IgG and IgM. The proteins were separated by their isoelectric points followed by size separation by SDS PAGE. Cross reacting spots from western Blots stained for IgG (A) and IgM (B) are mapped onto a gel stained with silver nitrate for protein. Blue marks indicate antigenic proteins that were detected for each. A total of 224 IgG reactive spots and 55 IgM reactive spots detected.

**S2 Fig.**
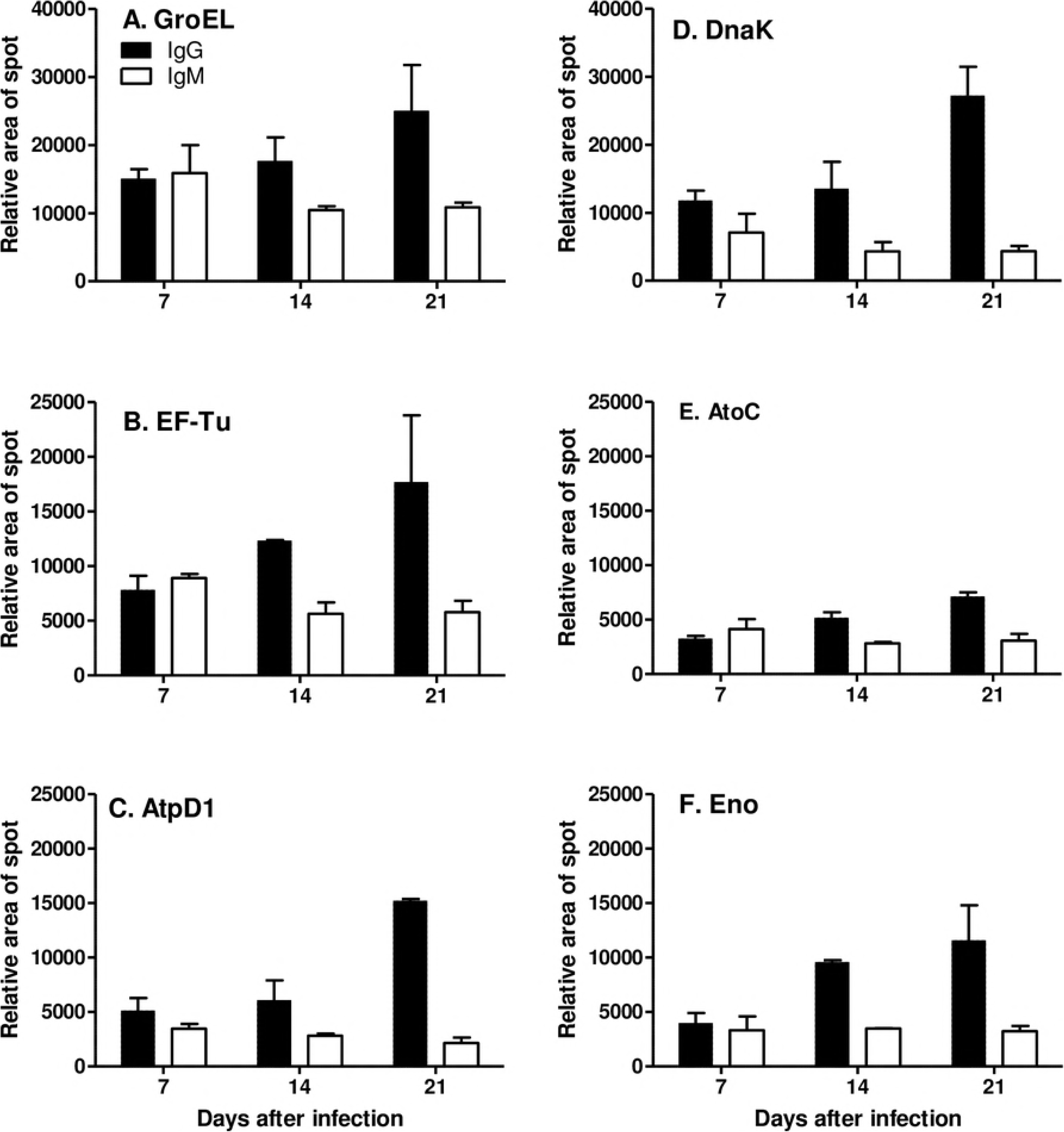
Characterization of goat humoral antibody responses to six proteins that were antigenic prior to *Burkholderia pseudomallei* strain MSHR511 infection. Goat humoral antibody responses of individual proteins of B. pseudomallei isolate (MSHR511). Goat sera were drawn prior to challenge or on days 7, 14 and 21 after infection (note: goat sera from day 16 were included with day 14 for calculations). The intensity of goat antibody response was calculated by western blot antigenic protein spot area. (A) Heat shock protein 60 family chaperone, GroL (B) Elongation factor Tu, EF-Tu (C) ATP synthase beta chain (D) Chaperone Protein DnaK (E) Sigmal-54 dependent DNA-binding response regulator, AtoC (F) Enolase, Eno. IgG (Black bars), IgM (White bars). Immunoglobulin G (IgG) ▬, and Immunoglobulin M (IgM) ▭

**S3 Fig.**
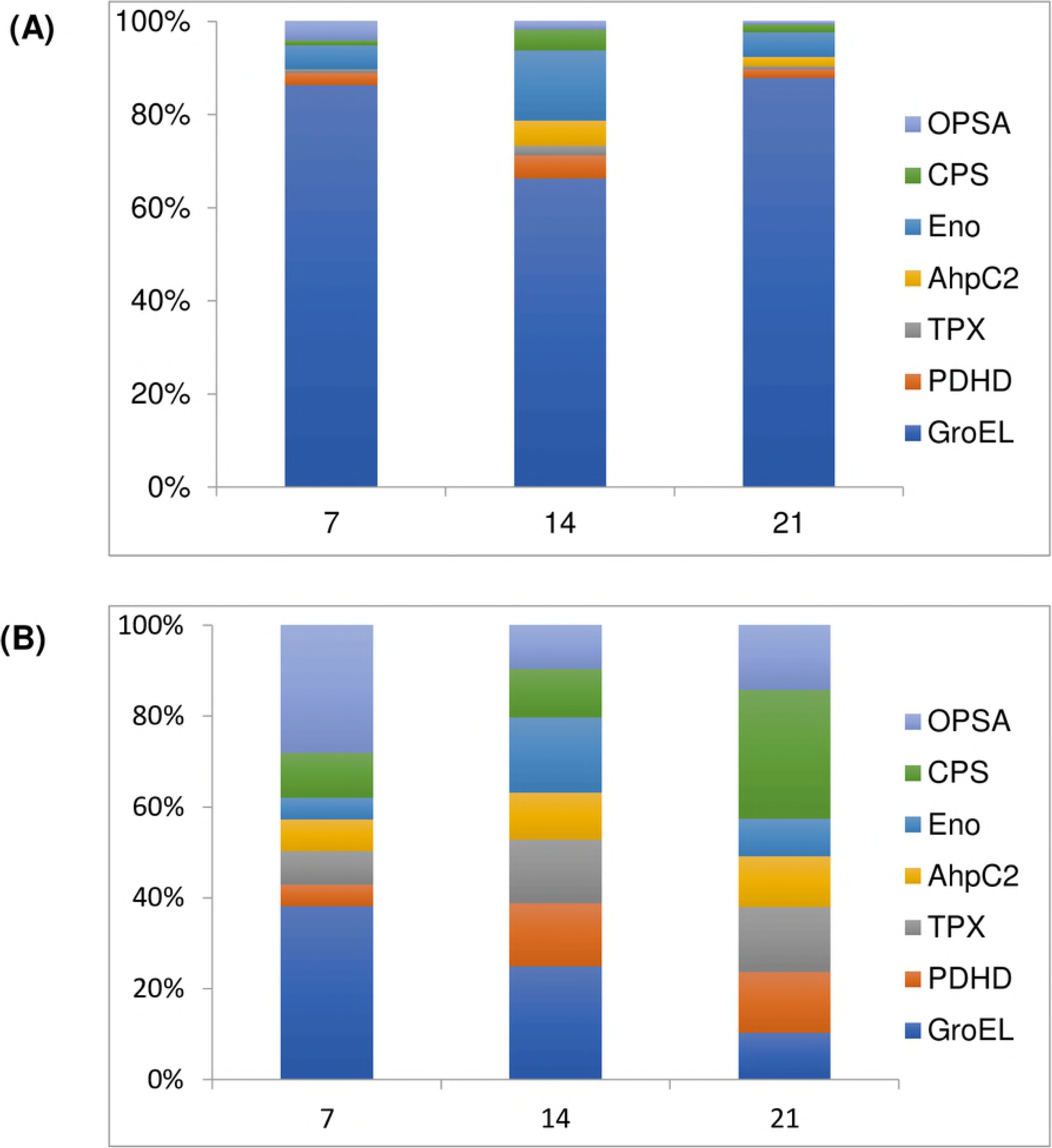
The relative contribution of seven individual antigens to the humoral antibody response. (A) Immunoglobulin IgG (IgG) antibody response was greatest for GroEL antigen for days 7, 14 and 21 followed by Eno on day 14. The humoral IgG response to the rest of the antigens (OPS A, CPS, AhpC2, TPX and PDHD) was typically <5% for each antigen. The high antibody response to GroEL antigen is thought may be due to memory B-cells being present in the immune circulation before challenge with *Burkholderia pseudomallei* strain MSHR511. (B) IgM antibody response was highest to GroEL antigen followed by OPS A for day 7 and these two antigens declined, respectively, for days 14 and 21. IgM response to CPS was relatively the same for days 7 and 14 but was most elevated compared to other antigens (Eno, AhpC2, TPX and PDHD) on day 21. The antibody response to the rest of the antigens (Eno, AhpC2, TPX and PDHD) was very similar.

S1 Table. Primers used to amplify the genes in this study S2 Table. List of purified antigens used in immunoassays

S3 Table. Antigenic proteins identified from an extract of *Burkholderia pseudomallei* using a non-redundant sequence database and MALDI-ToF mass spectrometry data.

S4 Table. Immunity (immune frequency) of antigenic proteins over days after infection.

